# Transcriptomic changes highly similar to Alzheimer’s disease are observed in a subpopulation of individuals during normal brain aging

**DOI:** 10.1101/2021.06.01.446628

**Authors:** Shouneng Peng, Lu Zeng, Jean-vianney Haure-mirande, Minghui Wang, Derek M. Huffman, Vahram Haroutunian, Michelle E. Ehrlich, Bin Zhang, Zhidong Tu

**Affiliations:** Institute of Genomics and Multiscale Biology, Icahn School of Medicine at Mount Sinai, One Gustave L. Levy Place, New York City, NY 10029, USA; Department of Genetics and Genomic Sciences, Icahn School of Medicine at Mount Sinai, One Gustave L. Levy Place, New York City, NY 10029, USA; Icahn Institute for Data Science and Genomic Technology, Icahn School of Medicine at Mount Sinai, One Gustave L. Levy Place, New York, NY 10029, USA; Departments of Medicine, Albert Einstein College of Medicine, 1300 Morris Park Avenue, New York, NY 10461, USA; Institute for Aging Research, Albert Einstein College of Medicine, 1300 Morris Park Avenue, New York, NY 10461, USA; Departments of Molecular Pharmacology, Medicine and Institute for Aging Research Albert Einstein College of Medicine, 1300 Morris Park Avenue, New York, NY 10461, USA; Departments of Psychiatry and Neuroscience, Icahn School of Medicine at Mount Sinai, New York City, NY, USA; Mental Illness Research, Education and Clinical Center (MIRECC), James J. Peters VA Medical Center, Bronx, NY, USA; Department of Neurology, Icahn School of Medicine at Mount Sinai, New York, NY, USA; Departments of Pediatrics and Genetics and Genomic Sciences, Icahn School of Medicine at Mount Sinai, New York, NY, USA

**Author notes:** Corresponding author: Zhidong Tu, IMI 3-70F, 1425 Madison Ave, New York City, NY 10029.

**Keywords:** Aging, LOAD, gene expression, brain transcriptome, brain aging subgroups, hippocampus, brain regions, Alzheimer’s disease

## Abstract

Aging is a major risk factor for late-onset Alzheimer’s disease (LOAD). How aging contributes to the development of LOAD remains elusive. In this study, we examined multiple large-scale transcriptomic data from both normal aging and LOAD brains to understand the molecular interconnection between aging and LOAD. We found that shared gene expression changes between aging and LOAD are mostly seen in the hippocampal and several cortical regions. In the hippocampus, the expression of phosphoprotein, alternative splicing and cytoskeleton genes are commonly changed in both aging and AD, while synapse, ion transport, and synaptic vesicle genes are commonly down-regulated. Aging-specific changes are associated with acetylation and methylation, while LOAD-specific changes are more related to glycoprotein (both up- and down-regulations), inflammatory response (up-regulation), myelin sheath and lipoprotein (down-regulation). We also found that normal aging brain transcriptomes from relatively young donors (45-70 years old) clustered into several subgroups and some subgroups showed gene expression changes highly similar to those seen in LOAD brains. Using brain transcriptomic data from another cohort of older individuals (> 70 years), we found that samples from cognitively normal older individuals clustered with the “healthy aging” subgroup while AD samples mainly clustered with the “AD similar” subgroups. This may imply that individuals in the healthy aging subgroup will likely remain cognitively normal when they become older and vice versa. In summary, our results suggest that on the transcriptome level, aging and LOAD have strong interconnections in some brain regions in a subpopulation of cognitively normal aging individuals. This supports the theory that the initiation of LOAD occurs decades earlier than the manifestation of clinical phenotype and it may be essential to closely study the “normal brain aging” to identify the very early molecular events that may lead to LOAD development.

## Introduction

Alzheimer’s disease (AD) is the most common cause of dementia and about 6.2 million Americans live with the disease based on the Alzheimer’s Association 2021 report (2021). Aging is the major risk factor for late-onset AD (LOAD), which occurs at age 65 or older (Caselli et al., 2009) and represents over 95% of the AD cases. A study analyzing 1,246 subjects aged 30–95 years found that the risk of developing AD dramatically increases in APOE ε4 carriers who are 70 years or older (Jack et al., 2015). It has been well-recognized that normal brain aging and LOAD share multiple common features, e.g., aging brains often manifest certain degrees of cognitive impairment, memory loss, metabolic disturbances, bioenergetic deficits, and inflammation. Even though aging increases the risk of AD and the two processes share similarities in multiple aspects, a detailed brain-region-specific view of their interconnection at the molecular level is not fully available. It is also unclear which aging mechanisms are playing major contributions to AD development and why some individuals may age without major cognitive deficits while others develop AD (Koivisto et al., 1995; Herrup, 2010). To help address these issues, we performed a global comparison of the transcriptomes from normal aging and LOAD brains across multiple regions in a hope to gain new insights into the molecular interconnection between aging and AD.

Despite the fact that many transcriptomic studies have been performed to investigate aging and LOAD independently, only a few have compared normal aging and LOAD transcriptomic data in a systematic way (Cribbs et al., 2012; Berchtold et al., 2013; Mastroeni et al., 2017; Lanke et al., 2018). For example, Berchtold et al. used microarray to profile 81 aging and AD brains and found that synapse-related genes showed progressive down-regulation in both aging and AD (Berchtold et al., 2013). Using the same dataset, Lanke et al. performed an integrated analysis on young, aging and AD brains by constructing co-expression network models, and found that modules associated with astrocytes, endothelial cells and microglial cells were up-regulated while modules associated with neurons, mitochondria and endoplasmic reticulum were down-regulated, and these modules significantly correlated with both AD and aging. All these studies greatly helped our understanding of the interconnections between aging and AD; however, the previous studies were limited in the brain regions examined and more importantly, they all treated aging and AD samples as uniform groups, and did not consider the possible heterogeneity in either aging or AD.

In this study, we compared the gene expression profiles of normal brain aging (age ≤ 70) with LOAD (age ≥ 60) (Yang et al., 2015; Hodes and Buckholtz, 2016) to understand the similarity and difference between aging and AD across multiple brain regions. We also considered brain aging subgroups and compared the aging subgroups with LOAD.

## Materials and Methods

### Data collection and pre-processing

We compiled and processed multiple large-scale human brain aging and AD gene expression datasets. We summarize and describe each dataset below and list all the data we studied in Table 1.

**Table 1.** List of brain transcriptomic datasets used for obtaining aging and AD gene signatures.

| Datasets | Gene List ID | List Details (Tissue, traits) | # of genes<br>(UP/DOWN) | # of protein<br>coding genes | Sample<br>Size |
| --- | --- | --- | --- | --- | --- |
| GTEx “Aging” sets (age ≤ 70) |  |  |  |  |  |
| GTEx | G_AMY | Amygdala | 50(19/31) | 46(19/27) | 46 |
| GTEx | G_BA24_AC | Anterior_cingulate_cortex (BA24) | 1460(473/987) | 1208(320/888) | 56 |
| GTEx | G_CAU | Caudate (basal_ganglia) | 77(50/27) | 55(34/21) | 75 |
| GTEx | G_CRBH | Cerebellar_Hemisphere | 322(138/184) | 277(108/169) | 73 |
| GTEx | G_CRBL | Cerebellum | 427(213/214) | 340(162/178) | 87 |
| GTEx | G_CTX | Cortex | 97(34/63) | 78(30/48) | 71 |
| GTEx | G_BA9 DLPFC | Frontal_Cortex (BA9) | 15(8/7) | 12(5/7) | 61 |
| GTEx | G_HIPP | Hippocampus | 1804(950/854) | 1403(705/698) | 57 |
| GTEx | G_HYPO | Hypothalamus | 2140(822/1318) | 1650(490/1160) | 53 |
| GTEx | G_NC | Nucleus_accumbens (basal_ganglia) | 11(6/5) | 9(5/4) | 69 |
| GTEx | G_PUTM | Putamen (basal_ganglia) | 18(13/5) | 17(13/4) | 67 |
| UK “Aging” sets (age ≤ 70) |  |  |  |  |  |
| UK | UK_FCTX | Frontal cortex (FCTX) | 170(50/120) | 145(42/103) | 94 |
| UK | UK_TCTX | Temporal cortex (TCTX) | 678(169/509) | 510(91/419) | 82 |
| UK | UK_OCTX | Occipital cortex (OCTX) | 54(30/24) | 44(23/21) | 95 |
| UK | UK_WHMT | Intralobular white matter (WHMT) | 464(267/197) | 373(204/169) | 93 |
| UK | UK_CRBL | Cerebellum (CRBL) | 186(101/85) | 156(87/69) | 92 |
| UK | UK_PUTM | Putamen (PUTM) | 65(30/35) | 52(22/30) | 95 |
| UK | UK_THAL | Thalamus (THAL) | 12(8/4) | 9(5/4) | 91 |
| UK | UK_HIPP | Hippocampus (HIPP) | 959(544/415) | 755(431/324) | 93 |
| Other “AD” sets |  |  |  |  |  |
| MSSM | MSSM_FP | BM10(MSSM_FP) | 54(20/34) | 47(19/28) | 135 |
| MSSM | MSSM_IFG | BM44(MSSM_IFG) | 71(24/47) | 61(24/37) | 116 |
| MSSM | MSSM_PHG | BM36(MSSM_PHG) | 1577(936/641) | 1391(868/523) | 103 |
| MSSM | MSSM_STG | BM22(MSSM_STG) | 269(150/119) | 242(142/100) | 122 |
| Mayo | Mayo_CRBL_Source * | Cerebellum | 2632(1360/1272) | 2282(1231/1051) | 151 |
| Mayo | Mayo_CRBL | Cerebellum | 1880(1008/872) | 1600(901/699) | 151 |
| Mayo | Mayo_TCTX_Source * | Temporal cortex | 3842(2034/1808) | 3437(1933/1504) | 151 |
| Mayo | Mayo_TCTX | Temporal cortex | 2951(1663/1288) | 2628(1578/1050) | 151 |
| ROSMAP | ROSMAP_DLPFC | Dorsolateral prefrontal cortex (DLPFC) | 162(89/73) | 141(80/61) | 241 |
| Annese2018 | Annese2018_HIPP | Hippocampus | 2122(808/1314) | 1925(742/1183) | 10 |
| Rooij2019 | Rooij2019_HIPP | Hippocampus | 2840(1109/1731) | 2667(1045/1622) | 28 |
| Jager | Jager_Clinical_AD | Clinical_AD in DLPFC | 855(466/389) | 740(412/328) | 478 |
| Jager | Jager_Cognitive_decline | Cognitive_decline in DLPFC | 3035(1481/1554) | 2662(1281/1381) | 478 |
| Jager | Jager_Tau_tangles | Tau_tangles in DLPFC | 238(155/83) | 209(133/76) | 478 |
| Jager | Jager_B_amyloid | B_amyloid in DLPFC | 2315(1158/1157) | 2020(1060/960) | 478 |
| Jager | Jager_Patho_AD | Patho_AD in DLPFC | 98(58/40) | 85(51/34) | 478 |
| MS “AD” sets |  |  |  |  |  |
| MS | MS_BM10_FP | Frontal Pole | 19(5/14) | 17(5/12) | 63 |
| MS | MS_BM17_OCTX | Occipital Visual Cortex | 345(325/21) | 297(279/19) | 53 |
| MS | MS_BM20_ITG | Inferior Temporal Gyrus | 397(397/0) | 349(349/0) | 58 |
| MS | MS_BM21_MTG | Middle Temporal Gyrus | 635(81/565) | 571(66/516) | 58 |
| MS | MS_BM22_STG | Superior Temporal Gyrus | 25(0/25) | 24(0/24) | 60 |
| MS | MS_BM23_PCC | Posterior Cingulate Cortex | 34(7/27) | 29(7/22) | 58 |
| MS | MS_BM32_AC | Anterior Cingulate | 239(33/207) | 214(30/184) | 59 |
| MS | MS_BM36_PHG | Parahippocampal Gyrus | 55(7/49) | 48(6/42) | 60 |
| MS | MS_BM38_TP | Temporal Pole | 44(28/16) | 39(25/14) | 58 |
| MS | MS_BM4_PCG | Precentral Gyrus | 131(8/123) | 122(7/115) | 49 |
| MS | MS_BM44_IFG | Inferior Frontal Gyrus | 874(589/566) | 764(513/495) | 53 |
| MS | MS_BM46_DLPFC | Dorsolateral Prefrontal Cortex | 58(11/47) | 53(8/45) | 57 |
| MS | MS_BM7_SPL | Superior Parietal Lobule | 346(240/110) | 314(220/97) | 50 |
| MS | MS_BM8_SFG | Superior Prefrontal Gyrus | 953(426/670) | 869(386/617) | 56 |
| MS | MS_CD | Caudate Nucleus | 31(30/1) | 29(28/1) | 52 |
| MS | MS_HIPP | Hippocampus | 107(25/82) | 95(22/73) | 55 |
| MS | MS_PUTM | Putamen | 94(52/42) | 82(45/37) | 52 |

\* AMP-AD Mayo gene signature is based on SourceDiagnosis

### GTEx brain data

GTEx brain gene expression data (v7 and v8) from 13 brain regions were downloaded from the GTEx (Genotype-Tissue Expression) portal (Consortium, 2015) and NIH dbGaP database. Donor ages ranged between 20-70 years and we removed samples from donors annotated with any brain diseases from further analysis. We corrected sex, collection center (batch), RIN (RNA Integrity Number), PMI (postmortem interval), and top 3 genotype principal components (PCs) to calculate gene expression associated with donors’ chronological age.

### UK brain data

Data from 134 post-mortem brain donors free of neurological conditions were obtained from the UKBEC dataset (UK Brain Expression Consortium) (Millar et al., 2007; Trabzuni et al., 2011). 10 brain regions were included, namely, cerebellum (CRBL, n = 131 samples), frontal cortex (FCTX, n = 128), hippocampus (HIPP, n = 123), medulla (specifically inferior olivary nucleus, MEDU, = 120), occipital cortex (specifically primary visual cortex, OCTX, n = 130), putamen (PUTM, n= 130), substantia nigra (SNIG, n= 102), temporal cortex (TCTX, n = 120), thalamus (THAL, n = 125) and intralobular white matter (WHMT, n = 132). To make the donor age distribution comparable between GTEx and UK, we only considered samples from donors whose age ≤ 70. The sample size for each brain region after filtering by age ≤ 70 is listed in Table 1.

### Mount Sinai brain data

364 human brains (238 females and 126 males) were accessed from the Mount Sinai/JJ Peters VA Medical Center Brain Bank (MSBB–Mount Sinai NIH Neurobiobank) cohort. These samples represented the full spectrum of cognitive and neuropathological disease severity in the absence of discernable non-AD neuropathology. Donor ages for the samples ranged 61–108. For microarray profiling of 19 brain regions, a linear model was adopted to identify genes differentially expressed among different disease stage groups using R package Limma with default parameters and corrected for covariates including sex, postmortem interval (PMI), pH, and race (Wang et al., 2016). For each brain regions, DEGs (Differential Expression Genes) with FDR ≤ 0.05 in any of the 6 traits (CDR, Braak, CERAD, PLQ_Mn, NPrSum, NTrSum in contrast of High vs. Low, Low vs. Normal or High vs. Normal) were combined as the DEGs for that brain region (see Table S1)(Wang et al., 2016).

### Other brain data

The AMP-AD knowledge portal (Hodes and Buckholtz, 2016) hosts datasets from three studies: Mayo Clinic Brain Bank (Mayo), the Religious Orders Study and Memory and Aging Project (ROSMAP) and Mount Sinai School of Medicine (MSSM). Mayo dataset included temporal cortex (80 AD, 71 controls) and cerebellum (79 AD, 72 controls). ROSMAP dataset contained samples from dorsolateral prefrontal cortex BA9 region (155 AD, 86 controls). The MSSM cohort included samples from four brain regions: parahippocampal gyrus, inferior frontal gyrus, superior temporal gyrus, and the frontal pole (n = 476, Table 1). DEGs were downloaded from the portal and filtered for genes with FDR ≤ 0.05.

### Jager AD gene signatures

De Jager et al. performed analyses on 478 ROSMAP dorsal lateral prefrontal cortex (DLPFC) tissue samples (Mostafavi et al., 2018). 5 gene lists were considered containing genes whose expression was associated with AD-related traits including clinical diagnosis of AD at the time of death, cognitive decline, tau, amyloid, and pathologic diagnosis of AD.

### Annse2018 and Rooij2019 AD gene expression signatures

Annse2018 profiled hippocampus CA1 gene expression in 10 males with age between 60 ∼ 81(5 AD and 5 controls) (Annese et al., 2018). DEGs with |log2(Fold Change)| > 1 and FDR ≤ 0.05 were selected. Rooij2019 profiled gene expression of hippocampus samples from 18 AD and 10 controls. DEGs with differential expression score ≥ 0.1 and FDR ≤ 0.05 were considered (van Rooij et al., 2019).

### Differential expression analysis and age-associated gene expression identification

Differential expression (DE) analysis in aging and AD was performed using the R package edgeR and Limma (Robinson et al., 2010; Law et al., 2014) and we adjusted batch, RIN, sex, and PMI in GTEx data and brain source, gender, batch effects and PMI in UK data. A linear regression model was applied to identify gene expression changes associated with age and we adjusted the same covariates as we did in the DE analysis (Yang et al., 2015; Zeng et al., 2020).

### Subgroup identification using hierarchical clustering

We used hierarchical clustering method to identify subgroups in normal aging GTEx and UK brain samples. We selected the top 5000 most variable genes which showed similar clustering result with using all genes (the adjusted Rand index > 0.9) for the hierarchical clustering of GTEx data. Ward.D2 method in the R hclust function was used (Murtagh and Legendre, 2014).

### Deconvolution of GTEx bulk tissue gene expression data to infer cell-type composition

The immunopanning-isolated cell RNAseq data covering 5 cell types: neuron, astrocyte, endothelial cell, microglia and oligodendrocyte in normal temporal lobe cortex was used as reference (Zhang et al., 2016) to infer the cell-type composition of GTEx hippocampal samples. Based on a recent work of deconvolution of GTEx brain samples across multiple regions (Patrick et al., 2020), we used the DSA (Digital Sorting Algorithm) (Zhong et al., 2013) for cell-type proportion estimation. We followed the recommended processing procedures (TMM normalization, top 100 markers) and applied DSA to HIPP and BA24_AC gene expression data (adjusted age, sex, PMI, RIN, batch, and 3 genotype PCs) to compare the cell-type proportions among GTEx aging subgroups and between brain regions.

### Comparison of PHG and GTEx hippocampal transcriptomes

We obtained gene expression data from 215 parahippocampal gyrus (PHG) samples which were profiled at Mount Sinai (Wang et al., 2018). To compare the PHG and GTEx hippocampal gene expression data, we first calculated log2(tmp + 1) for both datasets, merged these datasets and then removed the batch effects using R ComBat package with age, PMI, sex, RIN as covariates. Negative values were assigned to 0 followed by sample-wide quantile normalization. We selected the PHG samples with age > 70 and obtained 78 samples (19 normal (CDR = 0, braak score: bbscore ≤ 3, CERAD = “NL”), 59 LOAD (CDR ≥ 1, bbscore ≥ 5, CERAD = “definite AD”)). Based on the GTEx top 5000 variance genes, we clustered PHG samples and found the normal and AD samples were mixed to some degree. After removing the mixed samples, 51 samples (14 normal and 37 LOAD samples) showed clear separation into two groups corresponding to donors’ AD status. We also performed DE analysis in LOAD vs normal in filtered version (PHG 51 samples) and mixed version (PHG 78 samples) and compared the DEGs.

### Functional enrichment analysis

We annotated the biological functions of each gene list using DVAID tool (Huang et al., 2009; Sherman and Lempicki, 2009). We also performed pathway analysis using MetaCore integrated software suite (https://portal.genego.com/; last accessed in November 2019) to determine enriched biological processes. Signal transduction gene regulation network analysis was based on SIGNOR 2.0 (Licata et al., 2020) from the SIGnaling Network Open Resource (downloaded on Aug. 2020) using networkanalyst web tool (Zhou et al., 2019).

## Results

### 1. A global comparison of aging and AD signatures across brain regions

#### 1.1. Derive brain aging and AD signatures from multiple transcriptomic data

We collected multiple large-scale brain aging and AD gene expression datasets (Figure 1) which are summarized in Tables 1 and S1. It is of note that to obtain “normal” aging gene expression signatures, samples from donors annotated with any brain-related diseases were removed in the GTEx data. We used a linear regression model to identify gene expression associated with donors’ chronological age and consider these genes as brain aging genes (we call each list of brain aging genes as an aging signature for the corresponding brain region). We derived aging signatures in 11 out of 13 brain regions from the GTEx data; two regions, i.e., the brain spinal cord and substantia nigra showed no apparent age-associated genes (0 and 1 gene, respectively) and were not considered for further analysis. Similarly, we derived aging signatures from 8 out of the 10 brain regions profiled in UK data; substantia nigra and medulla tissues showed no apparent aging signatures and were removed from further analysis.

**Figure 1.**
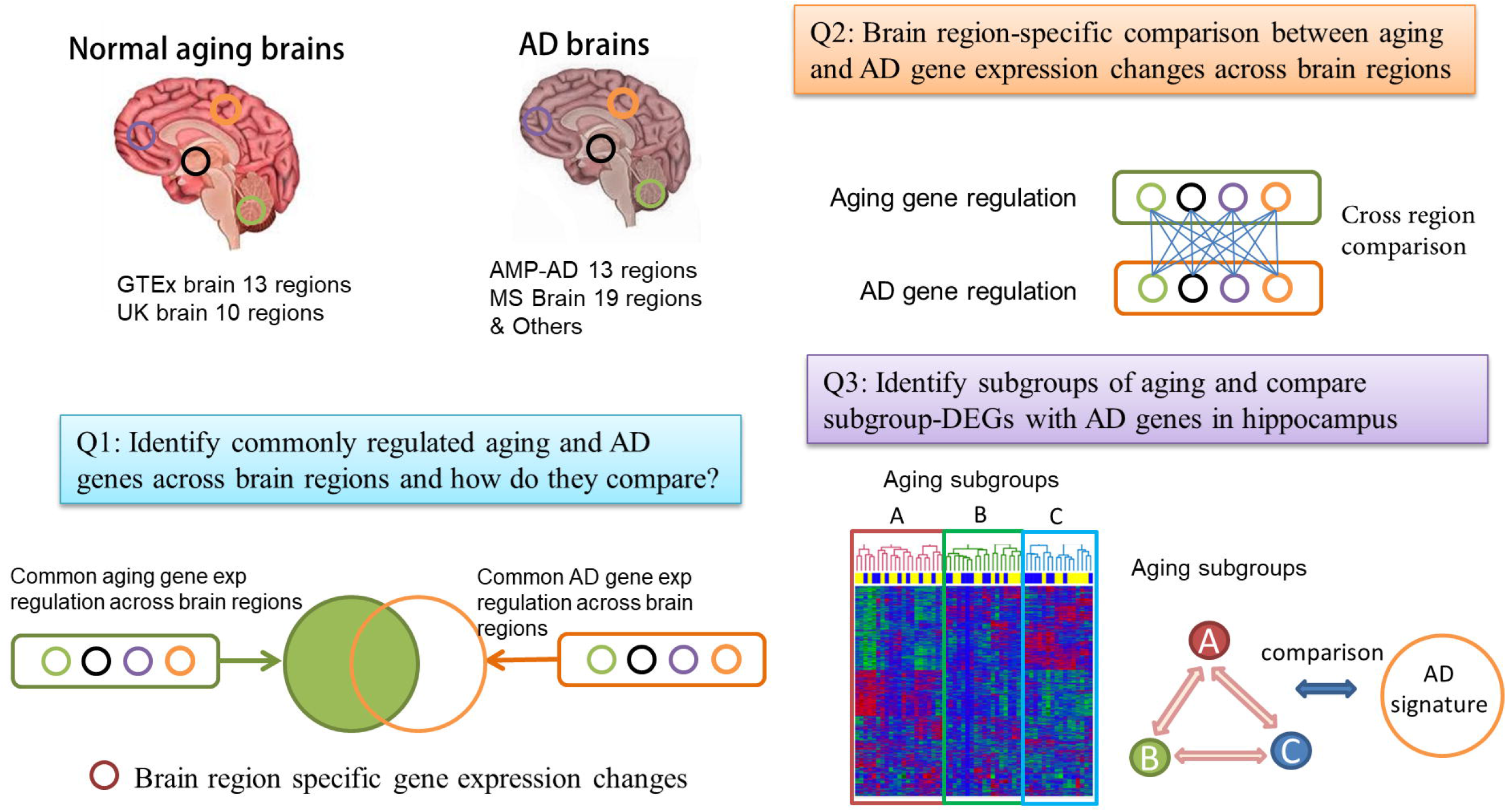
Flowchart of the comparison between normal brain aging and AD. We collect gene expression profiles from a large number of normal aging brain and AD brain samples across multiple brain regions. We perform both a global comparison (Q1) and region-specific comparison (Q2) of aging and AD transcriptomes. We also consider the subgrouping in aging brain samples and compare the aging subgroups with AD (Q3).

Genes differentially expressed between AD and normal control samples were extracted from previously published work (similarly we call them as AD signatures). These AD signatures are MS AD sets from Mount Sinai Medical Center Brain Bank (MSBB) AD cohort (Wang et al., 2016), and “Other” AD sets included AMP-AD knowledge portal data (Hodes and Buckholtz, 2016), Jager gene lists from the dorsal lateral prefrontal cortex (DLPFC) region (Mostafavi et al., 2018), Annese2018 DEG list from hippocampal (HIPP) CA1 (denoted as Annese2018 HIPP) (Annese et al., 2018) and Rooij2019 DEG list in HIPP (van Rooij et al., 2019) (denoted as Rooij2019 HIPP) (see Table 1 and Method).

#### 1.2. A global comparison of aging and AD signatures across brain regions

We first identified genes that show similar gene expression regulation across multiple brain regions in aging and AD datasets respectively (we call them global aging and AD signatures). We found that 91 aging genes are consistently up or down-regulated in more than 4 out of 19 aging gene lists (11 GTEx and 8 UK brain regions). Among them, 27 are up-regulated and 64 are down-regulated (see Table S2) with age. Similarly, 86 AD signature genes show consistent regulation in at least 9 out of 32 AD gene signature lists, among which 51 are consistently up-regulated and 35 are consistently down-regulated in AD brains (see Table S2). We then perform function annotation of these global aging and AD signatures.

##### 1.2.1. The global aging and AD signatures show up-regulation of immune complement pathway genes and down-regulation of synaptic related signaling genes

We used the DAVID tool to annotate the functional enrichment and found the 91 aging genes are enriched for synapse, nucleotide-binding and ion genes (see Table S3.1); while the 86 AD signature genes are enriched for synapse, phosphoprotein and transport genes (see Table S3.4). We further used MetaCore to annotate the function of consistently up- or down-regulated aging and AD signature genes. The 27 up-regulated aging genes are enriched for regulation of lipid metabolism, insulin regulation of glycogen metabolism (PHK gamma gene; FDR = 2.9E-2), immune/inflammation complement pathway (*C4B, C4, C4A*; FDR = 1.9E-06), response to metal ion/transport categories (e.g., cellular response to copper ion; FDR = 1.7E-06), and dopamine metabolic process (*MAOB, MAO*; FDR= 3.8E-3) (see Table S3.2); the 64 down-regulated aging genes are enriched for *MAPK* (FDR = 1.2E-02) / *ASK1* (FDR = 2.4E-02) pathway, synaptic vesicle / calcium ion exocytosis categories, synaptic related signaling (NMDA, glutamate) (see Table S3.3). The 51 up-regulated AD genes are involved in immune complement pathway (GPCRs), TGF-beta receptor signaling (FDR = 4.6E-02), cytoskeleton remodeling, and L-glutamate import/neurotransmitter transport (see Table S3.5); 35 down-regulated AD genes are involved in synapse and GABAergic neurotransmission, glutamate secretion/glutamatergic pathways, vesicle fusion/recycling and transport, cytoskeleton remodeling, cell adhesion and protein phosphorylation (see Table S3.6). Therefore, both AD and aging global signature genes show up-regulation in immune complement pathway and down-regulation in synapse (especially glutamate, synaptic vesicle recycling) related pathways; while aging genes are more involved in metal ion, transport, glycogen metabolism and AD signature genes are more involved in cytoskeleton remodeling, cell adhesion and synaptic categories.

##### 1.2.2. *SST* and *SVOP* are down-regulated and *FOXJ1, SLC44A1* are up-regulated in the global aging and AD signatures

Two genes, *SST* and *SVOP* are consistently down-regulated in both global aging and AD gene signatures, while *FOXJ1* and *SLC44A1* are consistently up-regulated.

*SST* (somatostatin) is a neuropeptide hormone that maintains permeability and integrity of the blood-brain barrier (BBB) by regulating *LRP1* and *RAGE* expression. It abrogates Aβ-induced JNK phosphorylation and expression of *MMP2* to maintain permeability and integrity of BBB (Paik et al., 2019). Reduction of SST levels in the CSF and brain tissue is associated with impaired cognitive function and memory loss (Solarski et al., 2018). *SVOP* (synaptic vesicle 2-related protein) can bind to adenine nucleotide (particularly NAD) via its C-terminal extremity(Yao and Bajjalieh, 2009). Although the functions of *SVOP* remain obscure, the evolutionary conservation and homology to transporters support it may play a role in molecular trafficking in synaptic vesicles(Janz et al., 1998).

In mammalian cells, *FOXJ1* is a member of the Forkhead/winged helix (FOX) family of transcription factors that is involved in ciliogenesis (Yu et al., 2008). It is shown to suppress *NFκB*, a key regulator in the immune response (Lin et al., 2004). It is hypothesized that *FOXJ1* may play a protective role involved in the pathophysiology of brain injury and may be required for the differentiation of the cells (Jacquet et al., 2009). The solute carrier 44A1 (*SLC44A1*) is a plasma membrane choline transporter. It is also a mitochondrial protein and acts as a choline transmembrane transporter. Choline is essential for the synthesis of the neurotransmitter ACh by cholinergic neurons for regulating neuronal activity. Choline deprivation in the central nervous system reduces acetylcholine (ACh) release, memory retention, and spatial cognition in the hippocampus (Canty and Zeisel, 1994; Nakamura et al., 2001; Zeisel, 2007).

### 2. Brain region-specific comparison of aging and AD signatures

In addition to comparing the global aging and AD signatures, we also compared aging and AD signatures in a brain region-specific manner.

#### 2.1. Hypothalamus, hippocampus, and certain cortical regions show stronger age-related gene expression changes compared to other brain regions

As can be seen in Figure S1, the number of aging genes varies across brain regions. The regions showing a large number of aging genes in the GTEx data are hypothalamus (HYPO: 490 UP and 1160 DN genes), hippocampus (HIPP: 705 UP and 698 DN genes), and brain anterior cingulate cortex BA24 (BA24_AC: 320 UP and 888 DN genes). For the UK data, the brain regions showing a large number of aging genes are hippocampus (HIPP:432 UP and 324 DN genes) and temporal cortex (TCTX: 91 UP and 419 DN genes). It is of note that GTEx and UK do not cover the same brain regions, e.g., the hypothalamus was profiled by GTEx but not in the UK brain data. Among the 4 regions profiled by both GTEx and UK, (i.e., HIPP, CRBL, PUTM, FCTX), HIPP and CRBL show a higher number of aging genes and stronger overlap between GTEx and UK aging signatures than the other two brain regions. Interestingly, the hippocampal aging signature significantly overlaps with aging signatures from several cortical regions, while the cerebellum aging signature has very little overlap with other regions. The cerebellum show the most distinguishable gene expression patterns from all other brain regions which was also reported by the Allen Brain Atlases study (Mahfouz et al., 2015).

Since a larger sample size provides greater statistical power and generally allows more age-associated genes to be identified, a fair comparison across brain regions requires the sample size to be identical or very close to each other. The GTEx HIPP (N = 57), BA24_AC (N=56) and HYPO (N=53) have relatively small sample sizes compared to other brain regions (see Table 1), so the larger number of age-associated genes in these brain regions were unlikely caused by their sample sizes. However, for the UK brain data, the hippocampal region had relatively large sample size (N=93) compared to other brain regions, such as the TCTX (N = 82), the brain region with the smallest sample size. To ensure the large number of age-associated genes identified in UK HIPP was not due to its sample size, we performed a down-sampling test in three brain regions to compare their age-associated genes when they have the same number of samples. As shown in Table S4, after down-sampling 565.76 ± 483.63 (mean ± standard deviation) age-associated genes could be identified in the UK HIPP. Although much fewer genes were found compared to the 959 age-associated genes identified from the 93 HIPP samples, the HIPP remains to show the largest number of aging genes among all the UK brain regions.

In summary, brain aging signatures are highly region-specific. Hippocampus, hypothalamus, and cortex TCTX and BA24_AC are more affected by aging on the transcriptome level than other brain regions surveyed by the GTEx and UK brain data even when the sample size difference is considered.

#### 2.2. Common hippocampal aging genes between GTEx and UK data

In general, the aging signatures from GTEx and UK show large reproducibility in matched brain regions. Using hippocampus as an example, the 705 up-regulated GTEx aging genes significantly overlap with 431 up-regulated UK aging genes by 77 genes (FDR = 2.6E-31); and the 698 down-regulated GTEx aging genes overlap with 324 down-regulated UK aging genes by 42 genes (FDR = 3.6E-12). The 119 commonly up- or down-regulated hippocampal aging genes between GTEx and UK are enriched for phosphoprotein, alternative splicing, acetylation, complement, and synapse (see Table S5.1). In addition to the immune / inflammation categories found in previous global aging signatures, the 77 commonly up-regulated aging genes are also enriched for *TGF-β* signal, transcription regulation (such as mRNA transcription by RNA polymerase II with FDR of 2.2E-03, ATP-dependent chromatin remodeling with FDR of 7.3E-03, blood vessel remodeling, membrane protein intracellular domain proteolysis, protein transport, and metabolic process (see Table S5.2). Furthermore, from signal transduction network analysis (see Table S5.4 and S5.5), the top signal network contains 67 up-regulated genes and is connected with many metabolic-related pathways such as endocrine resistance (FDR = 4.0E-15), *foxo* signaling pathway (FDR = 1.4E-13), thyroid hormone signaling pathway (FDR = 8.7E-12), sphingolipid signaling pathway (FDR = 1.8E-10), neurotrophin signaling pathway (FDR = 1.8E-10), autophagy – animal (FDR = 3.6E-10), and insulin resistance (FDR = 1.5E-09).

The 42 commonly down-regulated aging genes between GTEx and UK are related to insulin-like growth factor 1 (*IGF-1*) signaling, *ERK1/2* pathway (for caveolin-mediated endocytosis or signal transduction), methylation, cell adhesion (e.g., synaptic contact), and long-term potentiation pathways and previous reported energy-dependent pathways such as synaptic categories (e.g. synapse vesicle), oxidative categories (see Table S5.3) based on MetaCore analysis. *IGF-1* (insulin-like growth factor) is a growth factor and neurohormone with some evidence suggesting its involvement in neurocognitive functions, neuroinflammation, and amyloid-β clearance. Furthermore, from signal transduction network analysis on the KEGG database (see Tables S5.6 and S5.7), the top signal network is involved in circadian entrainment (FDR = 9.2E-07) and multiple synaptic-related pathways.

#### 2.3. Brain aging signatures from the hypothalamus, the hippocampus, and BA24 anterior cingulate cortex significantly overlap with AD signatures

After we evaluated the aging signatures between GTEx and UK data, we then compared the aging signatures with AD signatures in a brain-region specific manner. We further collected two hippocampal DEG lists (Rooij2019 and Annese2018, listed in “Other” AD sets) for the following analysis. As can be seen in Figure 2, strong overlap between aging and AD signatures are observed in hippocampus, hypothalamus, and several cortical regions. For example, the GTEx hippocampus aging signature strongly overlaps with AD signatures derived from either hippocampus or several cortex regions. For the 698 GTEx_HIPP_DN aging genes, they overlap with Rooj2019_HIPP_DN by 288 genes (adjusted P-value = 2.78E-130); and the 705 GTEx_HIPP_UP genes overlap with Mayo_TCTX _UP (1578) by 150 genes (adjusted P-value = 6.62E-28). It is of note that as we divide aging and AD signatures into up- and down-regulated genes, in most cases, the down-regulated aging genes only significantly overlap with down-regulated AD genes, and vice versa, which supports that these signatures represent real biological signals. The overlap pattern is highly brain region specific. For example, GTEx_HIPP_UP (705) significantly overlap with AD signatures from hippocampus, while it only overlaps with AD signatures from non-hippocampal regions such as MS_BM44_IFG_UP (513) by 21 genes (adjusted P-value = 0.72). In both GTEx and UK data, the aging signature in cerebellum shows relatively weak overlap with our AD signatures.

**Figure 2.**
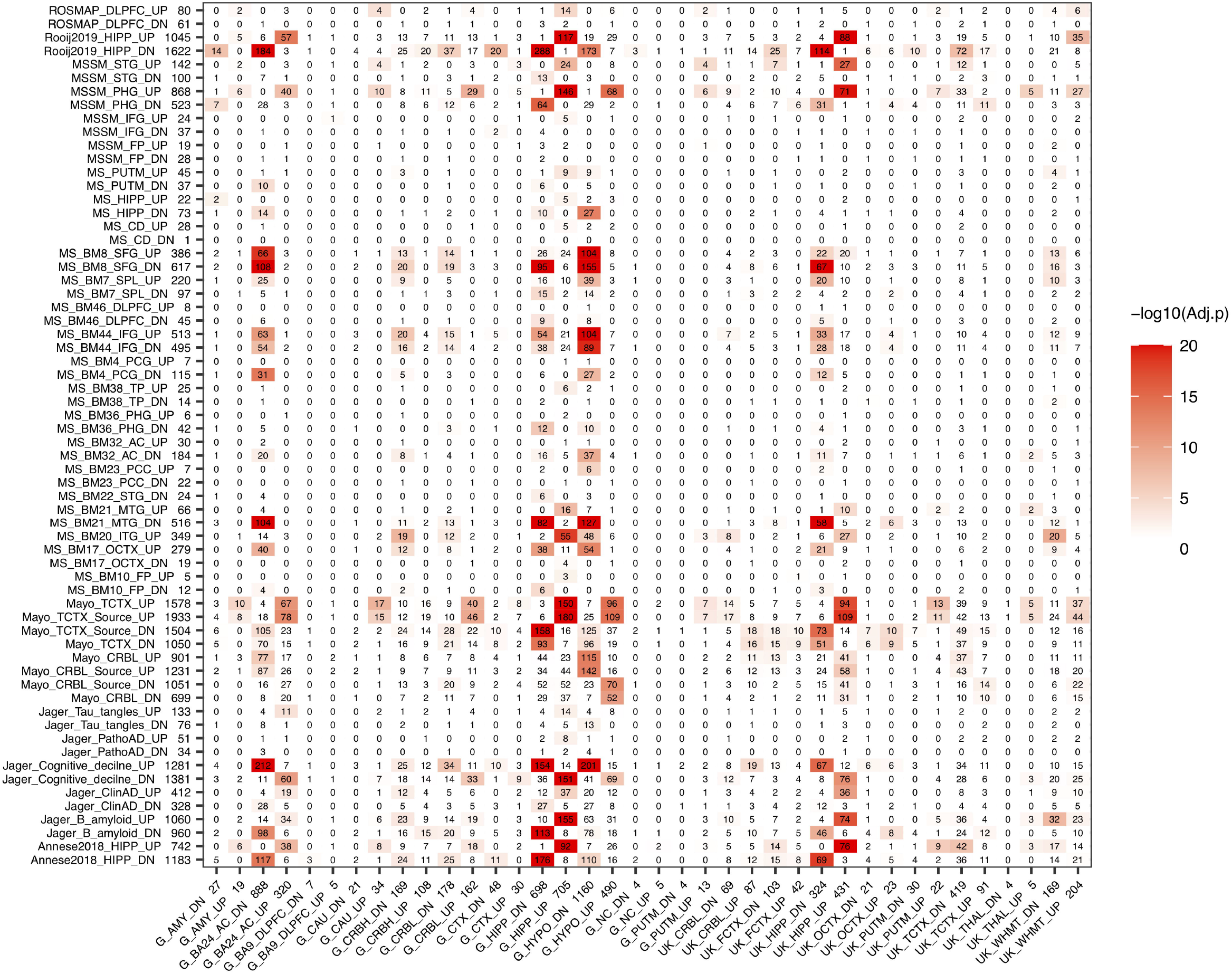
Overlap between aging and AD signatures in different brain regions. AD signatures are plotted in rows and aging signatures are plotted in columns. We separate each signature into up- and down-regulated genes and the number of genes in each signature is listed after its ID. The number in the heatmap indicates how many genes are common in the corresponding aging and AD signatures while the color indicates the significance of the overlap.

We also observed that the GTEx hippocampus and UK hippocampus aging signatures showed similar overlap pattern with AD signatures across different brain regions. Similarly, different AD signatures such as Annesse2018 and Rooj2019 showed similar overlap pattern with aging signatures across different brain regions, which further suggests that the aging and AD signatures from different studies are biologically meaningful and contain real signals of aging and AD.

#### 2.4. Functional enrichment of aging specific, aging/AD common and AD specific genes in hippocampus

Since aging and AD gene signatures show strong overlap in the hippocampus as seen in Figure 2, we focused on this brain region to investigate the functional enrichment of aging and AD genes. To achieve better robustness, we annotated hippocampus aging genes derived from both GTEx and UK with hippocampus AD genes derived from both Annese2018 and Rooij2019 (Figure 3). We divided genes in the aging and AD signatures into three categories: aging specific signature genes (denoted as “ASGs”), aging-AD common signature genes (denoted as “AADGs”), and AD specific signature genes (denoted as “ADSGs”). Phosphoprotein and alternative splicing are enriched in ASGs, AADGs and ADSGs (also in both up- and down-regulated genes); while glycosylation / glycoprotein, immune / inflammatory, stress response, mitochondria are more enriched in ADSGs; acetylation, nucleus, and Ubl conjugation are mostly enriched in ASGs (see Table S6.1). Down-regulated process of ASGs, AADGs, and ADSGs are all associated with membrane/cytoskeleton categories. Down-regulated AADGs are enriched for synapses (including cell junction, cholinergic/GABAergic/dopaminergic/glutamatergic synapse, AMPA etc.) and ion/transport (Figure 3 and Table S6); The comparison suggests that different from the normal aging process, the AD-specific and aging/AD commonly affected genes are more enriched for glycoprotein, inflammatory response, synapse and mitochondrial dysfunction, while the aging process are more related to the nucleus, acetylation, coiled coil, which are less directly related to neurodegenerative phenotype. The functional annotation of AADGs suggests that dysregulation in transcription regulation, energy metabolism, membrane remodeling, extracellular vesicles (EV) and synapse pathways have already initiated and developed to some degrees in normal brain aging even though these individuals remain cognitively normal, while other biological processes such as inflammation/immune response are further escalated in AD patients.

**Figure 3.**
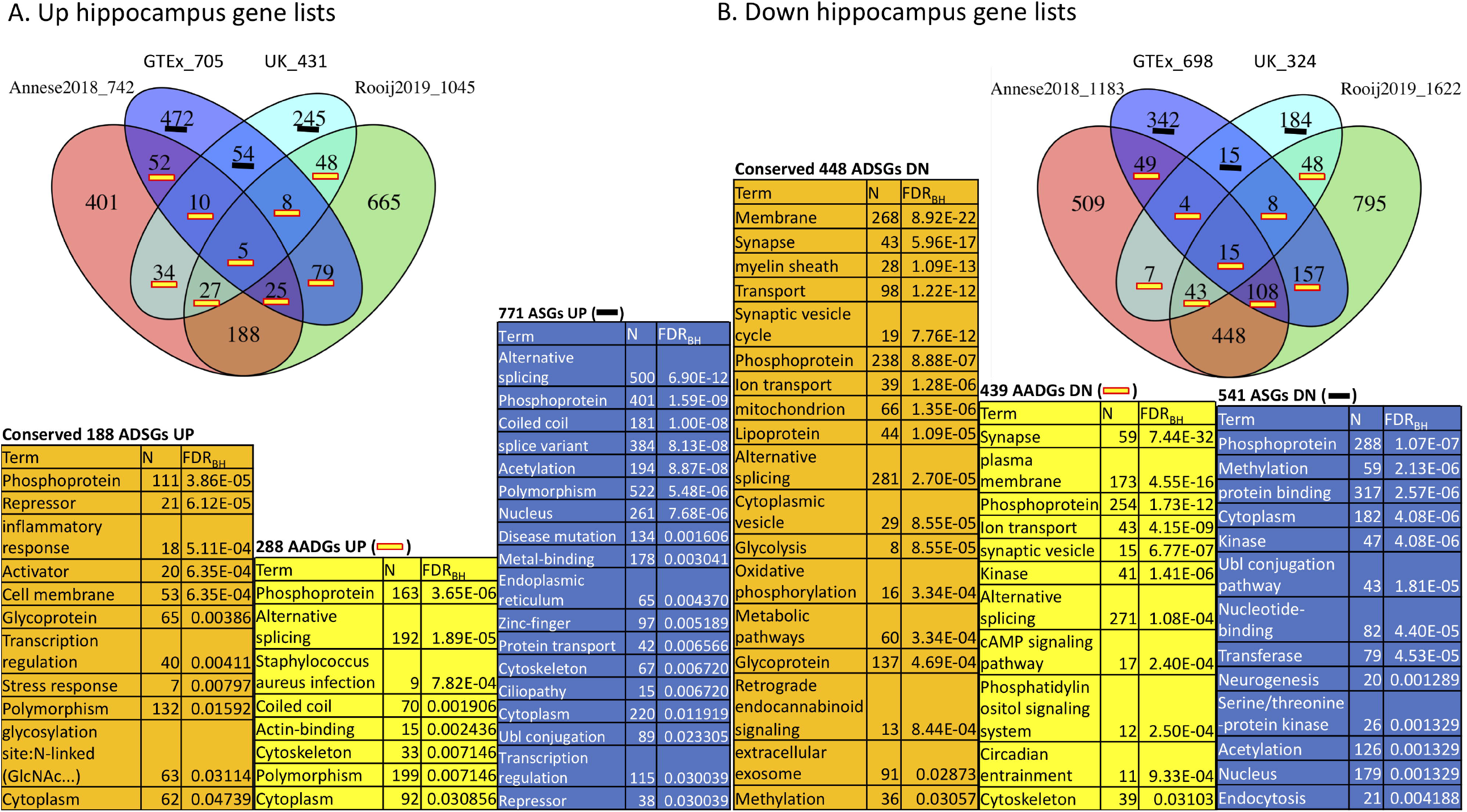
Function annotation of hippocampus aging signatures (GTEx, UK) compared with AD signatures (Annese2018, Rooij2019). We consider the functional enrichment of aging specific genes (ASGs), conserved AD specific genes (ADSGs) between two AD signatures, and the overlap between aging and AD genes (AADGs) which represents genes shared by at least one aging list and one AD list. We list the most representative function categories with FDR < 0.05. To reduce redundancy, only one representative functional category from each identified cluster of functions was selected.

### 3. Subgroup identification in normal aging brain hippocampus samples

#### 3.1 Subgroups can be found in both GTEx and UK hippocampus datasets

Although we have shown that aging and AD share multiple gene expression changes in the hippocampus, it is well known that not every old individual will develop AD. This suggests the interconnection between normal aging and AD may occur at different intensity within the aging population. To examine the potential heterogeneity of normal brain aging, we explored subgrouping in both GTEx and UK hippocampus samples; and we observed that both GTEx and UK samples can be divided into major subgroups. As shown in Figure 4, 56 GTEx hippocampal samples with donors’ age between 45 and 70 can form three major clusters (named subgroups A, B and C). We noticed that samples from different age groups are relatively evenly distributed across these subgroups, suggesting that subgrouping is not due to difference in donor ages. Similar subgrouping is also observed in UK hippocampal data (Figure 4B).

**Figure 4.**
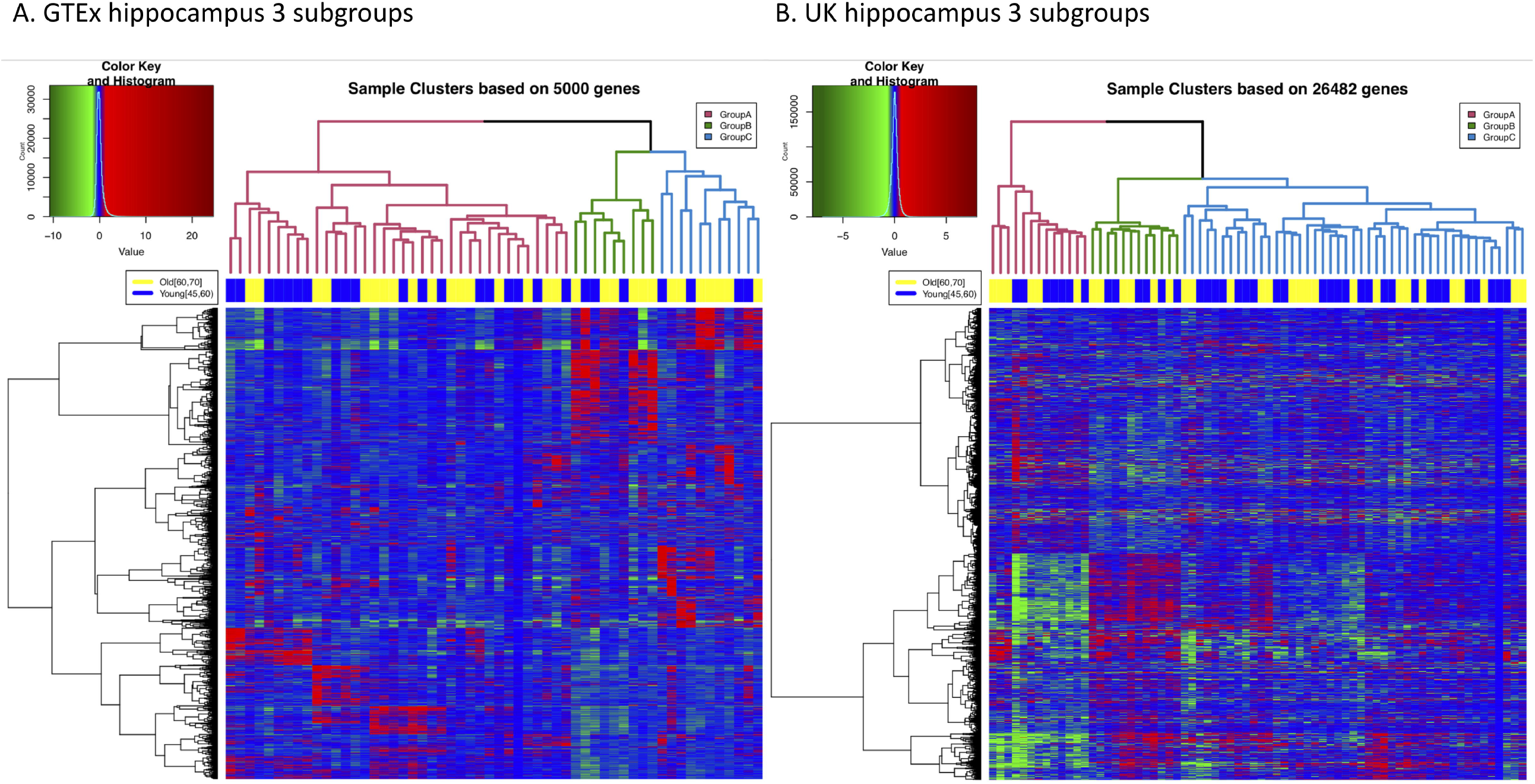
Three major subgroups can be identified from either GTEx or UK hippocampus gene expression data. (**A)**. Hierarchical clustering of GTEx hippocampus samples. 56 samples with donor age between 45 and 70 are plotted. Three subgroups are labeled in different colors in the dendrogram. The top color bar indicates that samples from different age groups are relatively evenly distributed into all the subgroups. (**B)**. Hierarchical clustering of 70 UK hippocampus samples (45 ≤ age ≤ 75) also suggests these samples can be divided into three major subgroups.

#### 3.2 DEGs derived from comparing aging brain subgroups highly overlap with AD signatures

We performed pair-wise comparisons to derive DEGs between subgroups in both GTEx and UK datasets, respectively. We jointly considered these DEGs (FDR < 0.01) with 3 hippocampal AD signatures (MS, Rooij2019, and Annese2018). As shown in Figure 5, DEGs derived from comparing GTEx subgroups B vs A (denoted as GTEx DEG BvsA) show strong overlap with AD signatures from Rooij2019 and Annese2018. For the GTEx_DEG_BvsA down-regulated genes (3018 genes), they overlap with Rooij2019 down-regulated genes (1622 genes) by 1274 genes while they only overlap with Rooij2019 up-regulated genes (1045 genes) by 7 genes. This is highly significant, as Rooij2019 down-regulated genes (1622 genes) overlap with another AD signature, Annese2018 down-regulated genes (1183 genes) by 614 genes, which is comparable to the overlap we see with aging subgroup DEGs. It is of note that GTEx samples used in the subgroup analysis are from donors 45-70 years old without AD or other types of brain diseases. Simply by performing subgrouping, the DEGs from comparing these subgroups highly overlap with AD signatures from independent studies, which further supports that gene expression changes in a subpopulation of the cognitively normal brains have very strong interconnection with AD. Similarly, GTEx DEGs CvsA also strongly overlap with AD signatures. Based on the pattern of overlap, we infer that GTEx subgroup A is more likely to be a “healthy” aging subgroup while GTEx subgroups B and C are more similar to AD (here denoted as “AD similar group”). Since GTEx DEG CvsB showed reverse overlap with AD signatures from Annese2018 and Rooij2019, this indicates that subgroup B is more similarity with AD samples compared to subgroup C. Therefore, the order of subgroups judged by how similar they are with AD samples can be inferred as AD samples > subgroup B > subgroup C > subgroup A, which is also consistent with the observation that GTEx DEG CvsA overlap with GTEx DEG BvsA in the same gene regulation direction. Similarly, for the UK data, we also observed very strong overlap between DEGs from comparing various subgroups and AD signatures. For example, the UK_DEG_AvsC_DN (3144 genes) highly overlap with Rooij2019_DN signature (1622 genes) by 762 genes. Similarly, we inferred the order of UK subgroups judged by how similar they are with AD signatures as AD > UK subgroup A >> UK subgroup B > UK subgroup C.

**Figure 5.**
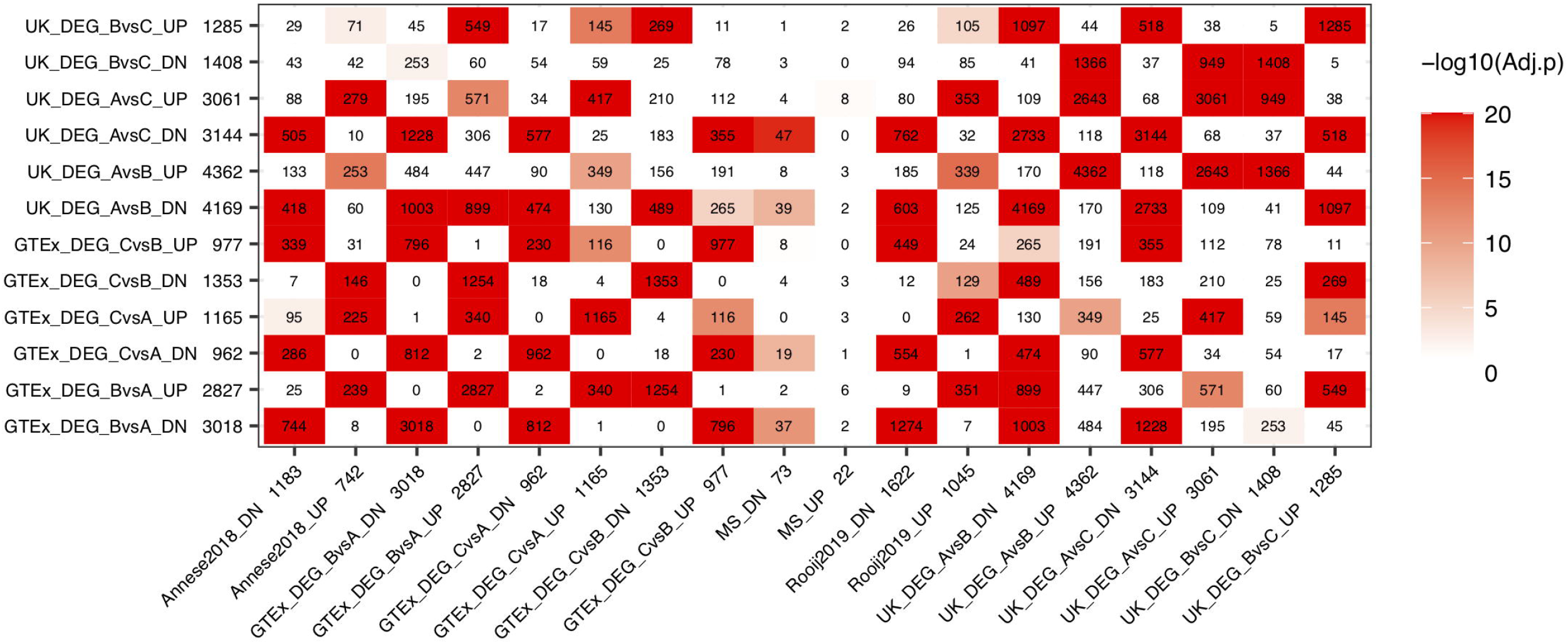
Comparison of DEGs between GTEx subgroups and AD signatures from hippocampus. “GTEx DEG CvsB” represents DEGs derived from comparing subgroups C vs. B in GTEx hippocampus, similar naming format is used for the rest gene lists.

#### 3.3 Function annotation of GTEx and UK subgroup DEGs with AD signatures

We next used the DAVID tool to annotate the functional enrichment for GTEx and UK subgroup DEGs (Figure 6, Table S7). We focused on the DEGs (FDR < 0.01) derived from comparing GTEx subgroup B (a subgroup that is more similar to AD) with the relatively healthy aging subgroup A (named GTEx DEGs BvsA) and similarly for UK DEGs AvsC with two AD signatures (Annese2018 and Rooij2019). Similar to previous annotations, we denote the aging subgroup DEGs as ABGs and annotate the functional enrichment of ABSGs (aging subgroup DEG specific genes), ADSGs, and ABADGs (common genes between aging subgroup DEGs and AD signature genes). As shown in Table S7.1 and Figure 6, certain post translational modifications (PTMs) (e.g., phosphoprotein, alternative splicing, Ubl conjugation, acetylation), membrane and cytoskeleton categories are significantly enriched in ABSGs, ADSGs, ABADGs and also enriched in both up- and down-regulated genes, respectively. ADSGs (both up- and down-regulation) are enriched for glycoprotein, while up-regulated ADSGs are more enriched for immunity, inflammatory response categories and down-regulated ADSGs are enriched for calcium/ion categories. The up-regulated conserved ABSGs show enrichment in cell cycle category and down-regulated conserved ABSGs show enrichment in proteostasis such as proteasome, protein transport and protein binding. For ABADGs, up-regulated genes are mainly enriched for transcription regulation and metabolism categories (e.g. PI3K-Akt signaling pathway, MAPK signaling pathway, insulin resistance), while down-regulated genes are mainly enriched for synapse, lipoprotein, and circadian entrainment categories.

**Figure 6.**
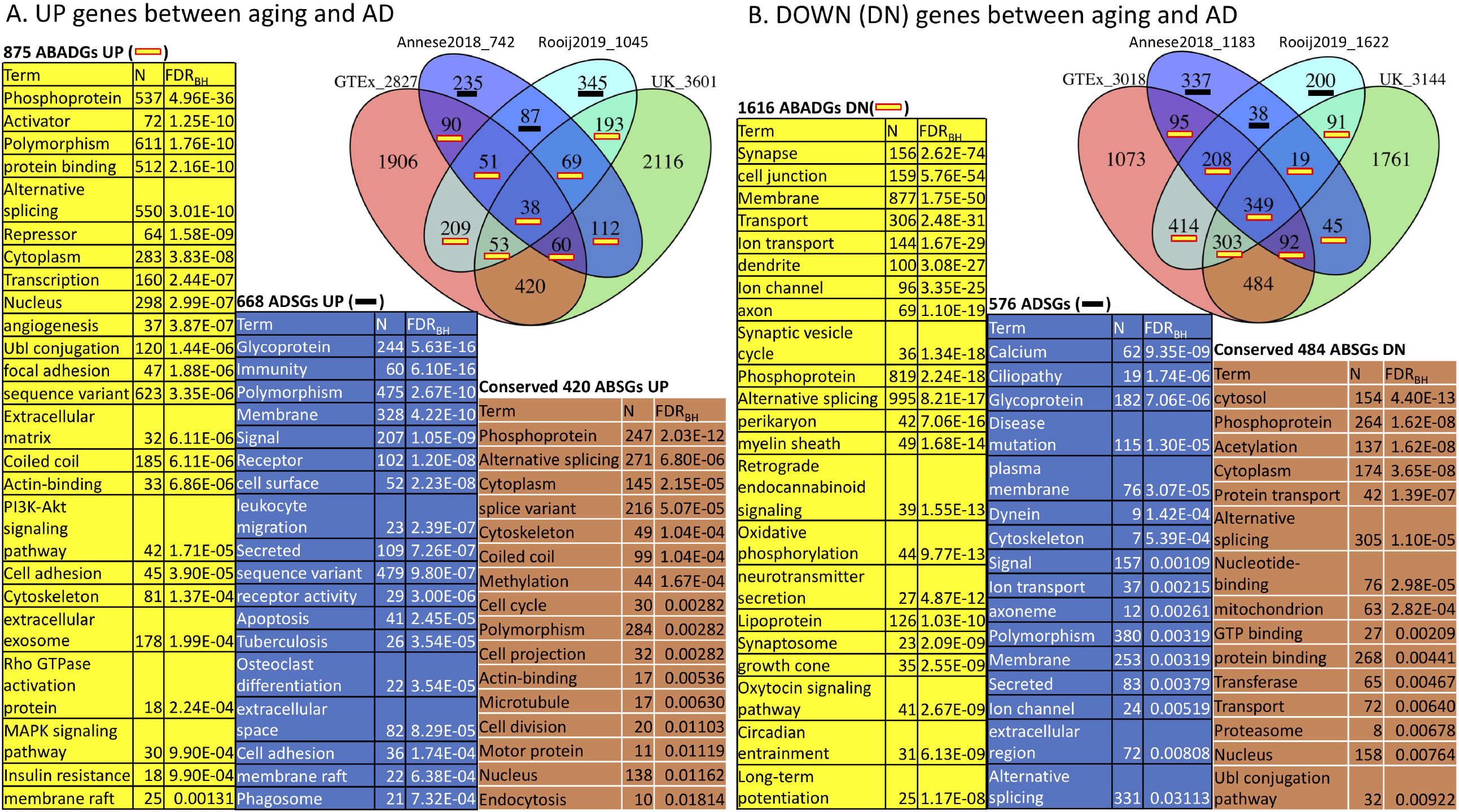
Function annotation of ABSGs, ABADGs, and ADSGs for a comparison of aging subgroup DEGs and AD signatures. The overlap of ABSGs between GTEx and UK is denoted as “Conserved ABSGs”. The union of Rooij2019 and Annese2018 subtracting any ABGs is denoted as “ADSGs”. We list the most representative function categories with FDR < 0.05. To reduce redundancy, only one representative functional category from each identified cluster of functions was selected.

In summary, aging subgroup DEG functional analysis further suggests that remodeling of PTMs (e.g., phosphoprotein and alternative splicing), proteostasis, cytoskeleton, and metabolism are likely initiated in normal aging subgroups that show highly similar gene expression changes with AD patients.

#### 3.4 Differentially expressed genes between subgroups could be partially driven by difference in the cellular compositions

The differentially expressed genes we observed between subgroups could be caused by either the dysregulation of gene expressions or the change in the cellular compositions or both. Since both GTEx and UK were bulk tissue gene expression data, the cellular composition information was not available. To evaluate if the cellular compositions were different between subgroups, we used computational tools to perform cell deconvolution. Based on a recent cell deconvolution work which provided experimental data validation (Patrick et al., 2020), we used the DSA (Digital Sorting Algorithm) method (Zhong et al., 2013) and cell-type reference data (Zhang et al., 2016) for cell-type proportion estimation (see methods). We were able to reproduce Patrick et al.’s result in the GTEx hippocampus when we made no adjustment to GTEx gene expression data (Fig S2A). When we adjusted the GTEx data by covariates like age, sex, PMI, we did notice some difference in the estimated proportion of several cell-types compared to Patrick et al.’s results (Fig S2B). When we applied this approach to GTEx HIPP gene expression data, as can be seen in Figure 7, neurons, oligodendrocytes and microglia showed significant difference among GTEx aging subgroups. Interestingly, the GTEx subgroup A (the heathy aging subgroup) is estimated to have higher proportions of neurons than AD similar subgroups; GTEx subgroup C (an AD similar group) showed to have elevated proportion of microglial cells; while GTEx subgroup B (another AD similar subgroup) showed to have elevated number of oligodendrocytes. Similarly, as shown in Figure S3, almost all the five cell-types’ proportions were different between hippocampus and BA24 (the two brain regions we chose to compare), suggesting cell-proportion difference widely existed across brain aging subtypes and brain regions.

**Figure 7.**
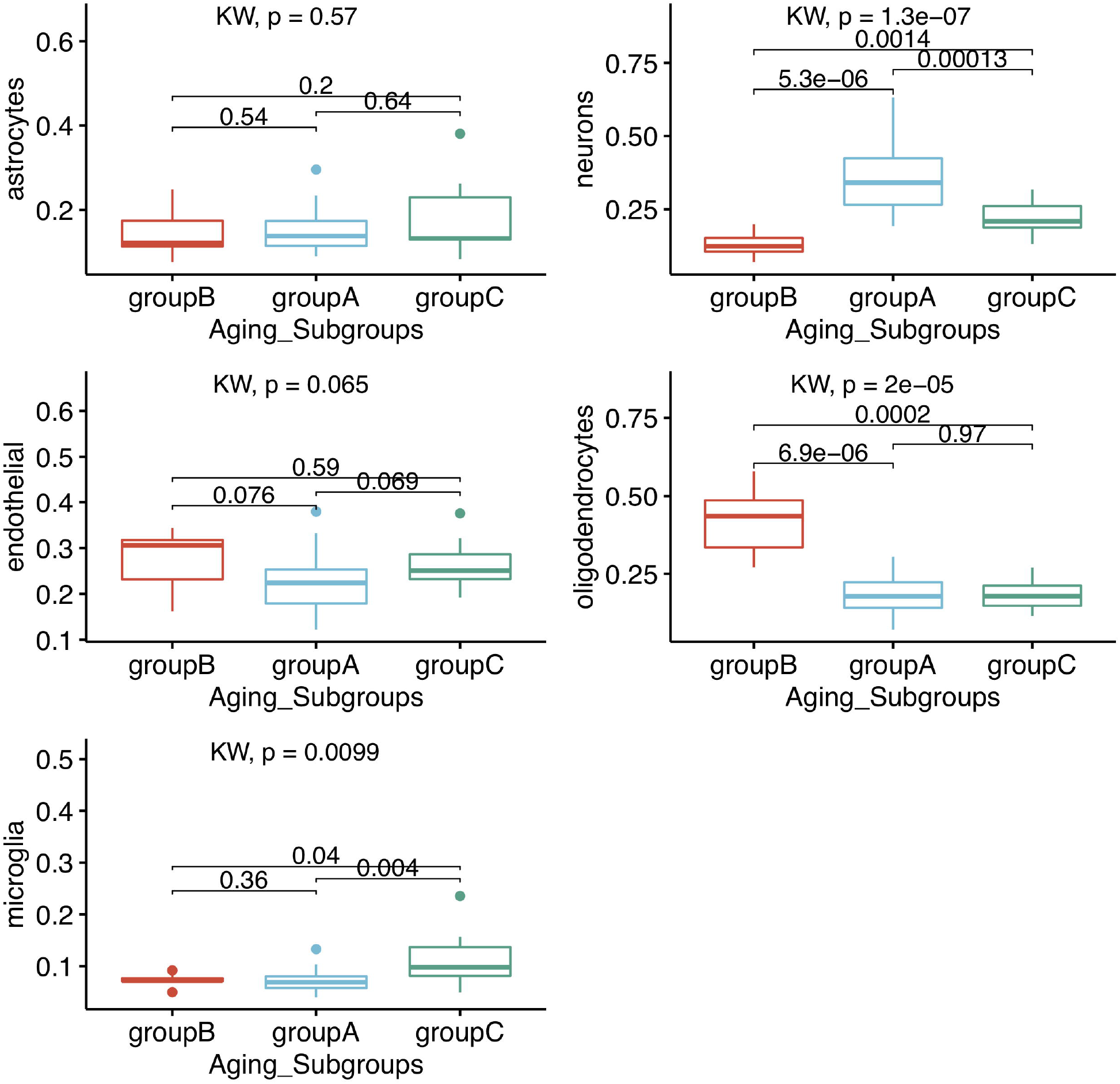
Subgroup comparison of estimated cell-type proportions of GTEx HIPP data using DSA method and Zhang’s reference data. Kruskal-Wallis rank sum test and wilcox.test rank sum test were used to calculate the significance levels between the groups.

#### 3.5 GTEx “healthy aging” subgroup will likely remain cognitively normal in older ages as implied from joint analysis with another brain transcriptomic dataset

In both GTEx and UK datasets, we observed a subgroup that is quite different from AD and subgroups that are more similar to AD on the transcriptome level. We hypothesize that the subgroup least similar to AD is a “healthy aging” subgroup and donors in this subgroup will have a better chance of being cognitively normal if they could live into 70s or older. In contrast, the “AD similar” subgroups will likely have a higher chance of developing cognitive deficits if these individuals could live into 70s or older. To test this hypothesis, a longitudinal study that follows-up these subgroup individuals for several decades will be required. This is not feasible mainly because brain tissues are mostly available from postmortem donors which do not permit longitudinal studies. To evaluate the hypothesis, we have to rely on alternative approaches.

To indirectly test the hypothesis, we relied on another large brain transcriptomic dataset which profiled more than 200 brain tissues from the parahippocampal gyrus (PHG) region (Wang et al., 2018). Although PHG is a different brain region, it is next to the hippocampus and our analysis showed that the two transcriptomic datasets are comparable. From these PHG samples, we selected two groups of samples, i.e., control and AD samples. The normal control samples were chosen from donors with age > 70, CDR = 0, braak score (bbscore) ≤ 3, CERAD = ‘NL’ which has 19 samples. For this group, the donor ages ranged from 73 to 103, with mean and standard deviation of 83.3 ± 8.7 (yrs). The second group is the AD group, which we required donors to be > 70 yrs, CDR ≥ 1, bbscore ≥ 5, CERAD = ‘definite AD’. 59 samples met these criteria, and the donor ages ranged from 71 to 104, with mean and standard deviation of 86.966 ± 8.4 (yrs). We merged these PHG data with the GTEx hippocampus RNA-seq data and corrected the batch effect using the R ComBat package (see method) to form a unified dataset. The joint processing introduced some changes to the GTEx gene expression. Therefore, we performed the hierarchical clustering on the 56 GTEx samples again and observed that the new subgrouping structure (subgroup B [n=11] > C [n=10] > A [n=35]) is highly similar to the previous clustering results (subgroup B’ [n=9] > C’ [n=11] > A’ [n=36]). For example, the new subgroup A (35 samples) overlaps with the previous subgroup A’ (36 samples) by 32 samples. We performed hierarchical clustering on the 78 PHG samples and found normal control and AD samples were partially mixed together (see Figure S4A). The partial mix of control and AD samples is not unexpected. For the control samples, they were from cognitively normal donors. Just like the GTEx and UK brain data, we have repeatedly observed that a subgroup of cognitively normal individuals showed gene expression changes highly similar to AD. On the other hand, there were “AD” samples mixed with the control samples, which is possibly due to the subtypes of AD samples(Neff et al., 2021) as in some AD subtypes, the PHG region could be relatively normal. Since we want to obtain samples that truly represent normal control and AD based on their gene expression, including the mixed samples will likely dilute the contrast between healthy and AD and blur the biological signals between the two. Based on this rationale, we removed the mixed samples and obtained 14 normal (ages 73 ∼ 103, 5 males and 9 females) and 37 AD samples (ages 73 ∼ 104, 11 males and 26 females) which formed two fully separated groups based on their gene expression (see Figure S4B). We then assigned these PHG samples to the GTEx subgroups by adding one PHG sample at a time to the 56 GTEx samples for hierarchical clustering. We found that all the 14 control samples clustered with the GTEx “healthy aging” subgroup A. For the 37 PHG AD samples, 5 (3 APOE ε3/ε4 and 2 APOE ε3/ε3) clustered with GTEx subgroup B and 26 clustered with GTEx subgroup C (the two “AD similar” subgroups), only 6 samples (all APOE ε3/ε3) clustered with the “healthy aging” subgroup A. This result suggests that the gene expression of the “healthy aging” subgroup is mainly associated with cognitively normal status in older individuals while the gene expression of the “AD similar” subgroup is largely associated with AD in this older cohort. To ensure the result is robust, we also compared the results when we kept the mixed sample and the results are summarized in Table S8. Without removing the mixed samples, for the 78 PHG samples (age > 70), 5 control samples were mixed with AD while 22 AD samples were mixed with controls (a total of 27 mixed samples). We assigned these PHG samples to the GTEx subgroups by adding one sample at a time using hierarchical clustering. 4 control samples clustered with GTEx group C and the other control sample clustered with GTEx group A. 7 AD samples clustered with GTEx group C, 1 clustered with group B (the first AD similar group), and 14 AD samples clustered with GTEx group A. This is largely consistent with our expectations, that the control samples mixed with AD clustered with an AD similar group, while majority of the AD samples that mixed with control samples (14/22) clustered with the healthy aging subgroup (GTEx subgroup A). Considering all the 78 PHG samples, our observation remains largely the same, i.e., most of the control samples (14 out of 19) clustered with the GTEx healthy aging subgroup, while majority of the AD samples (37 out of 59) clustered with GTEx AD similar subgroups. In addition, as shown in Figure S5, the filtered version (PHG 51 samples) captured much stronger gene expression differences between AD and control compared to the DEGs obtained from the mixed samples (PHG 78 samples).

## Discussion

To better understand the interconnection between aging and LOAD, we collected gene expression profiles from several large-scale transcriptomic datasets covering multiple brain regions and systematically compared aging and LOAD gene expression signatures. Different brain regions showed varied levels of gene expression changes in aging and AD, respectively. Among all the brain regions we studied, hippocampus is one of the top regions that show a very strong interconnection between aging and AD on the transcriptomic level. We observed common functional enrichment in aging and LOAD related to PTMs (especially for alternative splicing and phosphoprotein), neurotransmission (especially glutamate), membrane, cytoskeleton, and lipid metabolism. We also showed that gene expression changes in aging brain has more association with acetylation while AD-specific gene expression changes are more related to inflammatory response, glycoprotein, mitochondria and synapse. Importantly, we demonstrated that cognitively normal brains are not homogeneous in their transcriptomes and several major subgroups could be identified. By comparing gene expression among different subgroups, we showed that gene expression changes in a subpopulation have strong overlap with the AD signature, suggesting that although aging is a general risk factor for AD, only a subset of individuals may experience gene expression changes to a level that is significantly associated with the onset of the disease.

From our study, several biological processes shared between LOAD and normal aging could point to the mechanisms of how aging and AD interconnect. First, PTMs related genes such as alternative splicing and phosphoprotein are enriched in both up- and down-regulated ASGs, AADGs/ABADGs and ADSGs (e.g., top function terms in Figures 3 and 6). PTMs regulation plays critical role for synaptic plasticity at several levels (Kiltschewskij and Cairns, 2017) by increasing proteome diversity through alternative splicing, or by enabling activity-dependent regulation of mRNA localization, translation or degradation in the dendrite. It will be interesting to investigate what events trigger the PTMs that are observed in aging and AD. Second, the immune response strongly showed-up in AD signatures as can be seen in our results and previous studies (Verbitsky et al., 2004; Reichwald et al., 2009; Bordner et al., 2011; Cribbs et al., 2012). Since it was less significantly up-regulated in either ASGs/AADGs or ABSGs/ABADGs, this supports that the escalated inflammation is a hallmark of AD, relative to normal brain aging which is also known to be associated with low-grade neuroinflammation. Whether the inflammatory response plays a key causative role in the early stage of AD development or it is more of a reactive response to the upstream pathology requires further investigation.

Since it is infeasible to directly test if individuals in the “healthy aging” subgroup A will remain cognitively normal decades later, we relied on an alternative strategy in which we compared transcriptomes between GTEx brain aging subgroups with a cohort of older individuals who were either AD or cognitively normal. For the 51 PHG samples (14 control and 37 AD samples), we observed that 31 out of 37 AD samples clustered with GTEx subgroups B and C (AD similar subgroups); while all the 14 normal control samples clustered with GTEx subgroup A, which presumably represents a healthy brain aging group. This result suggests that the gene expression in subgroup A is strongly associated with normal cognitive functions in both relatively younger and older individuals, which supports that GTEx subgroup A is a healthy aging subgroup and individuals in this subgroup will likely remain cognitively normal when they get older. However, we did observe that several AD patients have their PHG transcriptome clustered with GTEx subgroup A (6 out of 37 AD samples). This indicates that an individual with a healthy aging gene expression pattern in the hippocampus or PHG can still have AD. We think this could be explained by the heterogeneity of AD which is increasingly known to have multiple subtypes (Neff et al., 2021). For example, at least four subtypes of AD (typical, limbic-predominant, hippocampal-sparing, and minimal atrophy AD) have been reported based on distribution of tau related pathology and regional brain atrophy (Ferreira et al., 2020). It is possible that some subtypes of AD could have their hippocampal transcriptomes similar to our healthy aging subgroup. For example, it has been reported that hippocampal-sparing AD subtype has a lower frequency of APOE ε4 compared with typical and limbic-predominant AD (Ferreira et al., 2020); interestingly, all the 6 AD samples clustered with the GTEx subgroup A are from donors of APOE ε3/ε3. Since AD is not a single brain-region disease, to accurately predict AD development based on gene expression or any type of brain region-specific data, we believe that multiple regions should be examined and studied together.

Although the PHG data suggest that different aging subgroups may have distinct probabilities of developing AD decades later, the “causal link” between certain aging subtypes and AD should not be assumed. Other alternative mechanisms should be considered which may explain our observations. For example, the aging subtypes could correspond to the natural fluctuation of brain states, while LOAD may represent a rather different state (or several states) that is difficult to escape once entered. To fully elucidate the underlying mechanisms of aging subtypes and their link to AD development, much more studies are needed.

The results from cell decomposition of the bulk gene expression data suggest that the differentially expressed gene expression between subtypes is at least partially due to the changes in cell proportions across the subgroups. For example, the down-regulation of synapse genes in the GTEx aging subgroup B might be explained by the decrease in the number of neurons in the samples in this subgroup. However, the deconvolution method we used assumed that the reference gene expressions do not change across conditions (in our case, the aging subgroups), which may not necessarily be true. Therefore, although it is very likely that cell compositions changed across subtypes, it is also possible that some gene expressions changed in a cell-type specific manner across the aging subgroups. To fully resolve this issue, single-cell profiling of samples from these aging subgroups will be highly useful.

Finally, we observed no significant difference in sex distribution among GTEx subgroups (Fisher exact test, p-value = 0.81) and UK subgroups (p-value = 0.37). For example, for the 36 samples in GTEx subgroup A, 27 were males, and 9 were females; for the 9 samples in GTEx subgroup B, 8 were males and 1 was female; and for GTEx subgroup C, 8 were males and 3 were females. Since there were only very limited number of female samples in the AD similar subgroups, we did not have sufficient statistical power to investigate the sex-related difference in these aging subgroups.

In summary, our study suggests that combined analysis of aging and LOAD can help us to understand how aging may contribute to the development of LOAD. Since most genomic studies on AD relied on AD samples from donors in 70s to 90s, these samples may not provide the information for the molecular events occurred in the very early stage of the disease development. As we demonstrated in this work, the brains from a subpopulation of cognitively normal individuals in their 40s-70s already show gene expression changes similar to AD, this supports that the initiation of LOAD could occur decades earlier than the manifestation of clinical phenotypes and it may be critical to closely study cognitively normal individuals in their 40s-60s to identify the triggering events in AD development.

## Acknowledgements

This work is supported by NIH grants R01AG055501, R01AG067312 to Z.T. and D.M.H. and R01AG057907 to B.Z. The content is solely the responsibility of the authors and does not represent the official views of the National Institutes of Health. This work was also supported in part through the computational resources and staff expertise provided by Scientific Computing at the Icahn School of Medicine at Mount Sinai.

## Author Contributions

ZT conceived the main concept of the work and BZ proposed the comparison with PHG data for “validation”. ZT and SP contributed to conceptualization, methodology, analysis, and preparation of the manuscript. SP, LZ, JH, MW, DH, VH, ME, BZ helped to discuss and improve the work. All authors have read and approved the manuscript for submission.

## Competing interests

No competing interests declared.

## Supporting Information

### Supplementary Figures

**Figure S1. Brain region specific overlap between GTEx and UK (age ≤70) aging signatures**.

**Figure S2. Estimated cell-type proportions of GTEx HIPP using DSA deconvolution method and Zhang’s cell-type reference data. (A)** GTEx HIPP without adjustment of covariates. Estimated cell-type proportions are highly similar to the results published by Patrick et al. **(B)** GTEx HIPP with adjustment of age, sex, PMI, RIN, batch, and 3 genotype PCs.

**Figure S3. Comparison of estimated cell-type proportions between GTEx HIPP and BA24_AC using DSA method and Zhang’s reference data**.

**Figure S4. Three major subgroups before and after mixed sample removal**. (A). Hierarchical clustering of 78 PHG samples with age > 70. Three subgroups are labeled in different colors in the dendrogram. The color bar on the top indicates from the AD status of the sample: normal or AD. **(B)**. Hierarchical clustering of 51 PHG samples with age > 70 after mixed samples being removed.

**Figure S5. Comparison of DEGs between LOAD vs. normal control samples from filtered version (PHG 51 samples) and mixed version (PHG 78 samples)**.

### Supplementary Tables

**Table S1**. Aging and AD protein coding filtered gene list

**Table S2**. Global Aging and AD signatures

**Table S3**. Function annotation for global Aging and AD genes

**Table S4**. Number of aging genes identified in 3 UK brain regions by down-sampling (DS) (DS to 81 samples were performed 50 times and aging genes were called using FDR <= 0.05).

**Table S5**. GTEx and UK hippocampus Aging genes function annotation

**Table S6**. Hippocampus Aging and AD genes by DAVID function annotation

**Table S7**. Hippocampus subgroups Aging and AD function annotation

**Table S8**. Subgroups in PHG and GTEx. GTEx subgroups ranked by their similarity to AD gene expression: B > C > A.

## Notes

### Competing Interest Statement

The authors have declared no competing interest.

### Summary of Updates

Updated the Section: Deconvolution of GTEx bulk tissue gene expression data to infer cell type composition

