## Supplemental Figures and Tables for "Transcriptomic changes highly similar to Alzheimer’s disease are observed in a subpopulation of individuals during normal brain aging"

**Figure S1. Brain region specific overlap between GTEx and UK (age ≤70) aging signatures.**

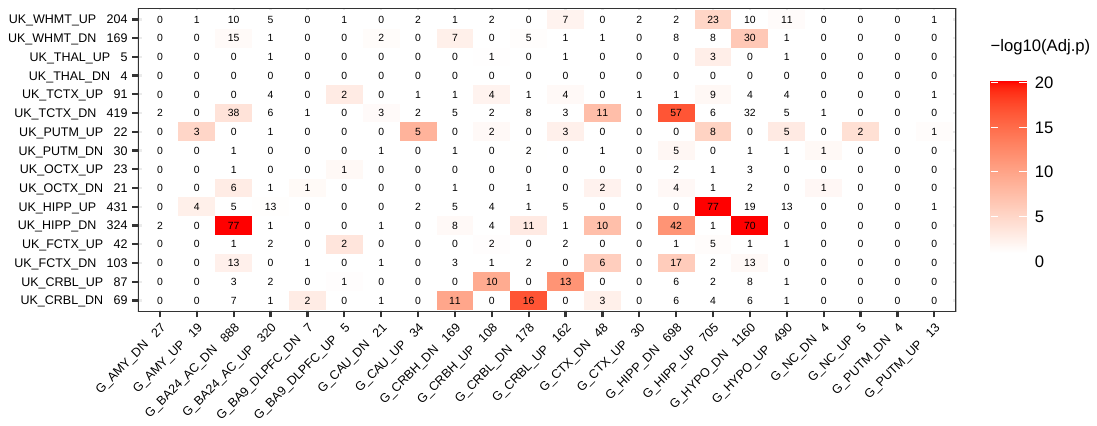

Notes: GTEx v7, signed as G; UK (age ≤70) signed as UK.

**Figure S2. Estimated cell-type proportions of GTEx HIPP using DSA deconvolution method and Zhang’s cell-type reference data. (A)** GTEx HIPP without adjustment of covariates. Estimated cell-type proportions are highly similar to the results published by Patrick et al. **(B)** GTEx HIPP with adjustment of age, sex, PMI, RIN, batch, and 3 genotype PCs. Astro: astrocytes, endo: endothelial cells, neuro: neurons, and oligo: oligodendrocytes.

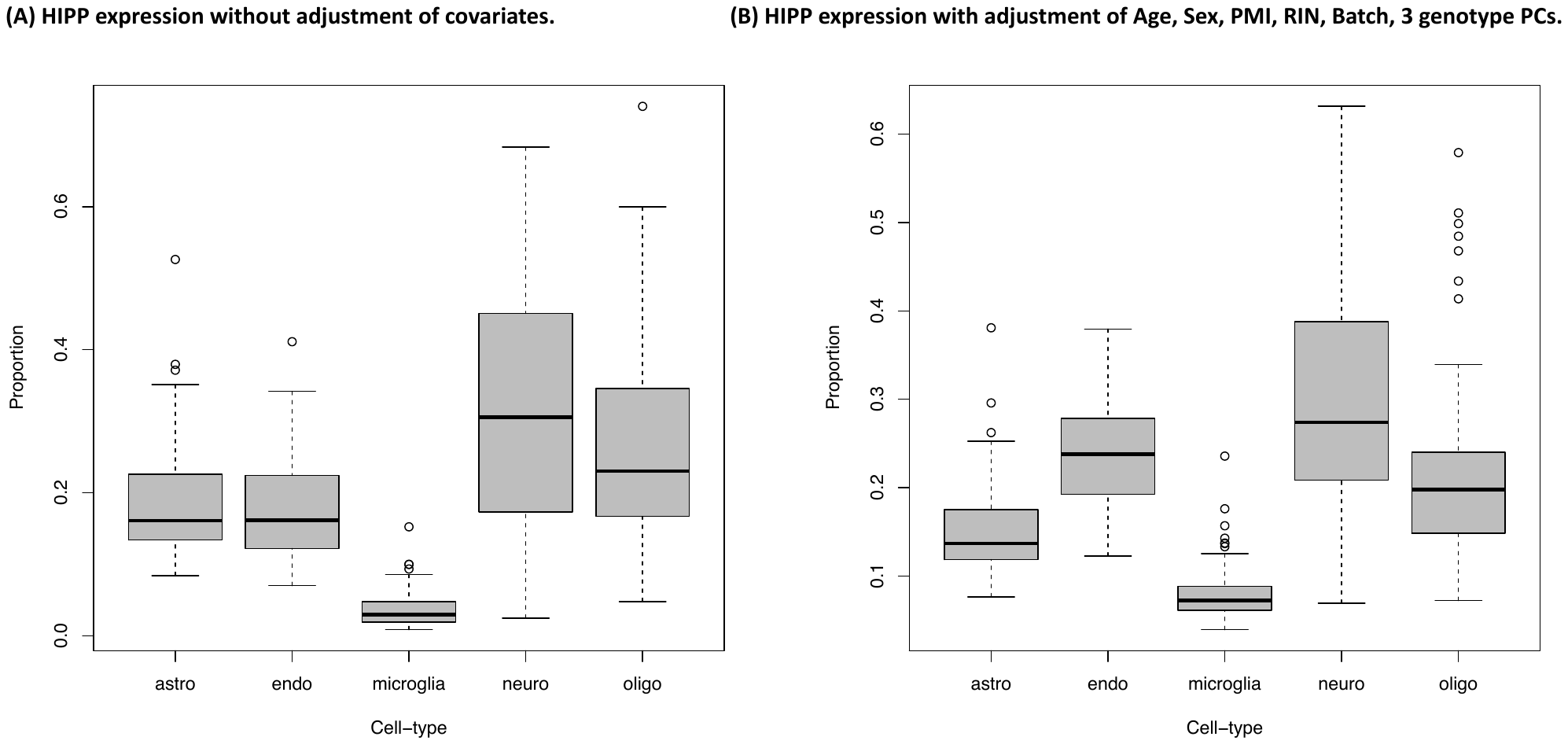

**Figure S3. Comparison of estimated cell-type proportions between GTEx HIPP and BA24_AC using DSA method and Zhang’s reference data.**

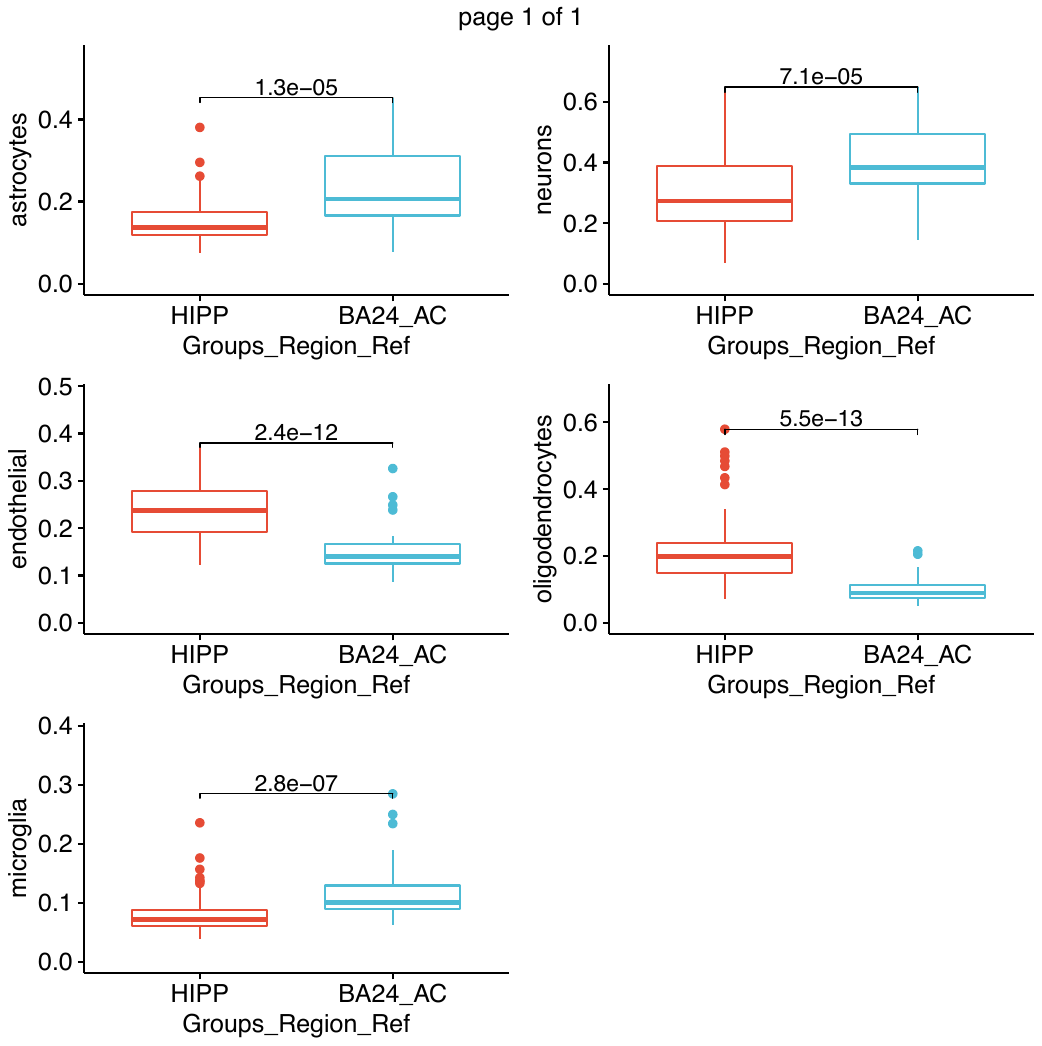

**Figure S4. Three major subgroups before and after mixed sample removal. (A).** Hierarchical clustering of 78 PHG samples with age > 70. Three subgroups are labeled in different colors in the dendrogram. The color bar on the top indicates from the AD status of the sample: normal or AD. (**B).** Hierarchical clustering of 51 PHG samples with age > 70 after mixed samples being removed.

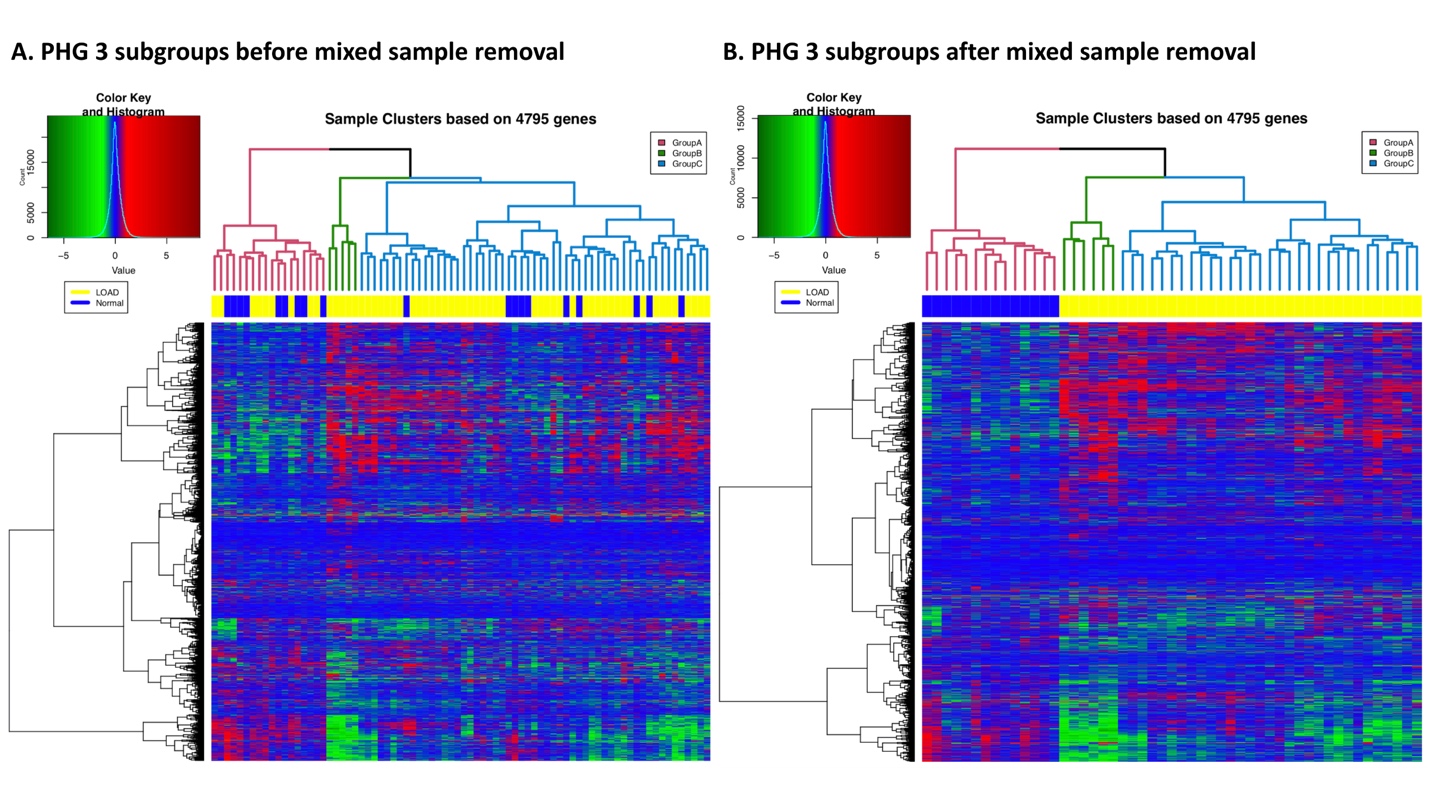

**Figure S5. Comparison of DEGs between LOAD vs. normal control samples from filtered version (PHG 51 samples) and mixed version (PHG 78 samples).**

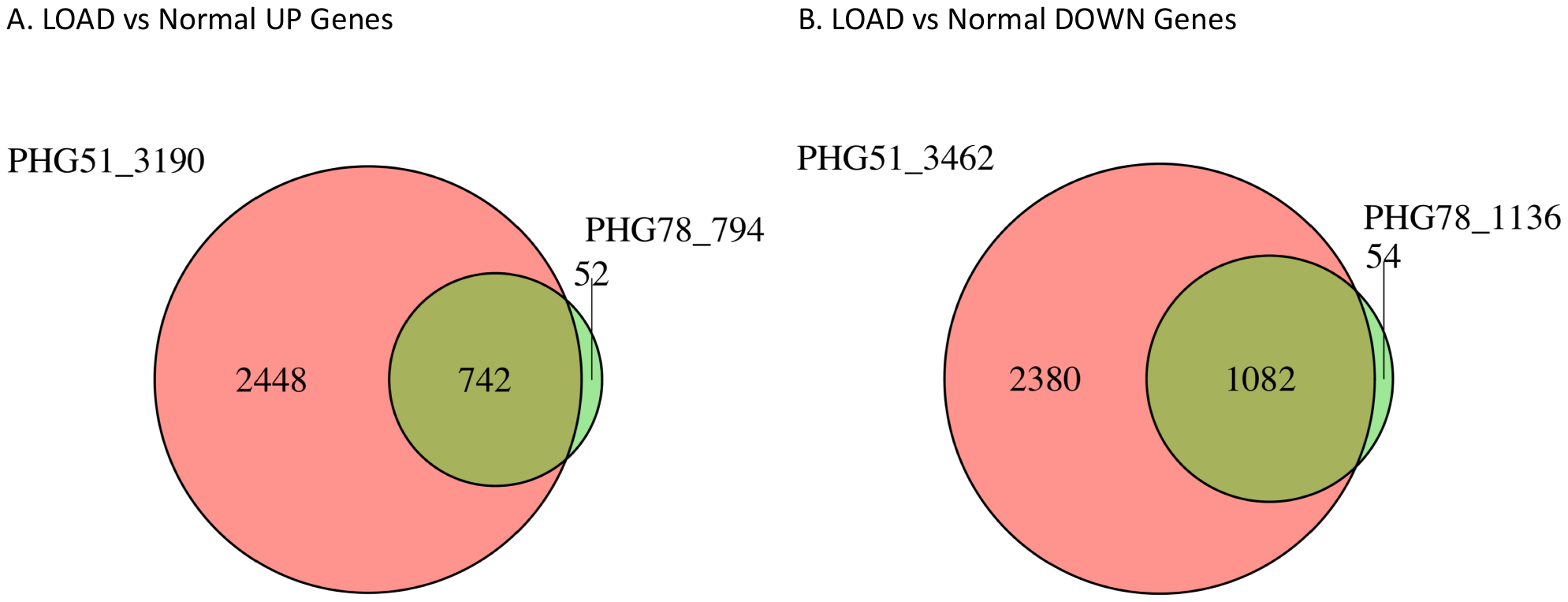

**Tables**

**Table S4.** Number of aging genes identified in 3 UK brain regions by down-sampling (DS) (DS to 81 samples were performed 50 times and aging genes were called using FDR <= 0.05).

|  | FCTX(N=94) | TCTX(N=82) | HIPP(N=93) |
| --- | --- | --- | --- |
| ALL Samples | 170 | 678 | 959 |
| 50 DS mean ± sd | 117.66 ± 66.13 | 538.2 ± 52.15 | 565.76 ± 483.63 |

**Table S8.** Subgroups in PHG and GTEx. GTEx subgroups ranked by their similarity to AD gene expression: B > C > A.

| No. of PHG samples clustered with GTEx subgroups (**Control** / AD) | Subgroup A  [“healthy”] | Subgroup B  [“AD similar”] | Subgroup C  [“AD similar”] | Total |
| --- | --- | --- | --- | --- |
| Filtered samples | 20(**14**/6) | 5(**0**/5) | 26(**0**/26) | 51(**14**/37) |
| Mixed samples | 35(**15**/20) | 6(**0**/6) | 37(**4**/33) | 78(**19**/59) |
| Removed samples | 15(**1**/14) | 1(**0**/1) | 11(**4**/7) | 27(**5**/22) |
